# IFNγ drives neuroinflammation and demyelination in a mouse model of multiple system atrophy

**DOI:** 10.1101/2022.11.22.517543

**Authors:** Nicole J. Gallups, Gabrielle M. Childers, Jhodi M. Webster, Asta Zane, Ya-Ting Yang, Nikhita Mudium, Fredric P. Manfredsson, Jeffrey H. Kordower, Ashley S. Harms

**Author notes:** Correspondence to: Ashley S. Harms, University of Alabama at Birmingham, 1719 6^th^ Ave South, Birmingham, Alabama 35294.

## Abstract

Multiple system atrophy (MSA) is a rare and fatal synucleinopathy characterized by insoluble alpha-synuclein (α-syn) cytoplasmic inclusions located within oligodendroglia. Neuroinflammation, demyelination, and neurodegeneration are correlated with areas of GCI pathology, however it is not known what specifically drives disease pathogenesis. Recently in a mouse model of MSA, CD4+ T cells have been shown to drive neuroinflammation and demyelination, however the mechanism by which this occurs also remains unclear. In this study we use genetic and pharmacological approaches in a novel model of MSA to show that the pro-inflammatory cytokine interferon gamma (IFNγ) drives neuroinflammation and demyelination. Furthermore, using an IFNγ reporter mouse, we found that infiltrating CD4+ T cells were the primary producers of IFNγ in response to α-syn overexpression in oligodendrocytes. Results from these studies indicate that IFNγ expression in CD4 T cells drives α-syn-mediated neuroinflammation and demyelination, and strategies to target IFNγ expression may be a potential disease modifying therapeutic strategy for MSA.

## Introduction

Multiple system atrophy (MSA) is a rare, rapidly progressive, and fatal demyelinating synucleinopathy with no known disease modifying therapy^1,2^. Unlike other synucleinopathies such as Parkinson disease (PD), MSA pathology is characterized by severe autonomic failure and rapidly progressive demyelination and neurodegeneration associated with alpha-synuclein (α-syn) containing glial cytoplasmic inclusions (GCIs) within oligodendrocytes^1,3,4^. MSA is divided into two subtypes, MSA-Parkinsonian (MSA-P) and MSA-Cerebellar (MSA-C). MSA-P is the most common, affecting 80% of patients and neurodegeneration occurs primarily within the striatonigral pathway^1^. MSA-C leads to olivopontocerebellar atrophy^1^. Although the areas of degeneration differ depending on MSA subtype, GCI pathology is present in vulnerable and resistant regions across the neuraxis. In addition to GCI’s within oligodendroglia, there is significant astrogliosis as well. GCI pathology is associated with significant neuroinflammation, demyelination, and neurodegeneration^1,2,5^. However, the mechanism by which GCI pathology leads to neuroinflammation, demyelination, and neurodegeneration is currently unknown.

Previous studies have shown neuroinflammation as a pathological hallmark of MSA. In MSA post-mortem brains, widespread astrogliosis and microgliosis are present within areas of neurodegeneration and demyelination^1,2,5^. The expression of major histocompatibility complex II (MHCII), the equivalent of human leukocyte antigen-DR (HLA-DR) in humans, is found on antigen presenting cells (APCs) within the CNS^6^. In post-mortem MSA brains, our previous study shows that α-syn GCI pathology is accompanied by MHCII+ expression and increased infiltration of peripheral T cells (CD4+, CD8+)^6^. Using a novel modified AAV in which human α-syn is overexpressed in oligodendroglia (Olig001-SYN)^7,8^ we observed significant neuroinflammation, demyelination, and neurodegeneration. Thus, effectively modeling MSA in rodents and non-human primates^9^. Using this Olig001-SYN model of MSA, we also demonstrated significant MHCII induction on CNS resident microglia and infiltrating macrophages, along with the infiltration of CD4 and CD8 T cells, similar to that observed in post-mortem brains^6^. Using mice that are genetically deficient in CD4 T cells, we found that Olig001-SYN induced-MHCII expression, infiltration of peripheral immune cells, and demyelination were attenuated, indicating a disease driving role of adaptive immunity, specifically CD4+ T cells in MSA pathogenesis^6^. Interestingly, upon further investigation of T cells isolated from Olig001-SYN transduced mice, α-syn overexpression resulted in significant production of the proinflammatory cytokine interferon gamma (IFNγ)^6^. As infiltrating T cells are a significant source of IFNγ, it is not currently known whether IFNγ mediates mechanisms that drive MSA pathogenesis.

IFNγ is an important pro-inflammatory cytokine. In response to proinflammatory stimuli, immune cells such as T cells, myeloid cells, natural killer cells, and B cells can produce IFNγ leading to MHC mediated antigen presentation^10,11^. IFNγ released from CD4+ T cells, particularly the Th1 subtype, enhances inflammation by binding to its receptor (IFNγR1) and, via the JAK/STAT pathway, activates genes responsible for T cell differentiation, T cell activation, and MHCII antigen presentation. In other demyelinating diseases like multiple sclerosis (MS), IFNγ is significantly increased. In experimental autoimmune encephalomyelitis (EAE), a mouse model for MS, neutralizing via an antibody against IFNγ attenuated neurodegeneration and neuroinflammation^12,13^. Furthermore, when Tbet (a transcription factor required for CD4+ T cell differentiation into Th1) was genetically knocked out in mice, it prevented the development of EAE^14^. Like MS, IFNγ is significantly increased in the CSF of MSA patients, however, no follow up studies have been conducted to determine if the IFNγ is pathogenic^15,16^. Although IFNγ is an important mediator of neuroinflammation in demyelinating disease^12,17^, there are currently no studies investigating a similar role of IFNγ in neuroinflammation and demyelination in MSA.

In this study, we sought to determine whether IFNγ is responsible for α-syn-mediated neuroinflammation and demyelination in the Olig001-SYN mouse model of MSA-P. Utilizing genetic and pharmacological approaches, global IFNγ targeting using genetic knockout mice or an IFNγ neutralizing antibody (XMG 1.2), attenuated α-syn mediated neuroinflammation and demyelination. Furthermore, using a novel Thy1.1/IFNγ reporter mouse, we determined that IFNγ was expressed primarily by CD4+ T cells and minimally in other immune cells in response to α-syn overexpression, suggesting that CD4 T cells mediates disease progression. These findings indicate that IFNγ, primarily produced by Th1 T cells, drives neuroinflammation and demyelination in MSA, highlighting IFNγ as a potential therapeutic target in the treatment of MSA.

## Materials and methods

### Mice

Male and female C57BL/6 (#000664 Jackson Laboratories) were used for these studies and maintained on a congenic background. IFNγ/Thy1.1 reporter mice with a C57BL/6 background (generously donated by Dr. Casey Weaver) were also used and have been previously described and characterized^18^. Additionally, male and female Tbet -/- mice (#004648 Jackson Laboratories) were used. Under a C57BL/6 background, these mice have exon 1 of the T-box 21 (*Tbx21*) deleted. All research conducted on animals were approved by the Institutional Animal Care and Use Committee at the University of Alabama at Birmingham (UAB).

### Olig001 vector

The Olig001 vector is a modified AAV capsid generated via directed evolution that has been characterized previously^7–9^. Briefly, the Olig001 capsid has a >95% tropism for oligodendrocytes and the vectors utilized contained the CBh promoter and bovine growth hormone polyA, controlling the expression of either transgene (human α-syn or GFP as control). The Olig001 vector was provided by the University of North Carolina Vector Core facility.

### Stereotaxic surgery

Male and female mice aged to 8-12 weeks, were anesthetized with isoflurane applied by an isoflurane vaporizing instrument provided by the Animal Resource Program at UAB. Using a Hamilton syringe and an automatic injecting system, mice were unilaterally (Luxol Fast Blue, DAB and immunofluorescence staining; n=5) or bilaterally (flow cytometry; n=5) injected with 2 μl of Olig001-GFP (1 ×10^13^ vector genomes (vg)/ml) or Olig001-SYN (1 ×10^13^ vg/ml) into the dorsolateral striatum at a rate of 0.5 μl/min to mimic MSA-P. The needle was left in the injection site for an additional 2 min and then slowly retracted over the course of 2 min. The stereotaxic coordinates used from bregma were AP + 0.7 mm, ML +/ – 2.0 mm, and DV – 2.9 mm from dura. All surgical protocols and aftercare were followed and approved by the Institutional Animal Care and Use Committee at the University of Alabama at Birmingham.

### Immunohistochemistry tissue preparation

Four weeks post transduction of the Olig001 virus, mice were anesthetized and transcardially perfused with 0.01M Phosphate-buffered saline (PBS) pH 7.4, followed by a fixation with 4% paraformaldehyde (in PBS, pH 7.4; PFA). Brains were dissected and incubated in 4% PFA solution for 4 hours at 4°C. After PFA fixation, the brains were cryoprotected in a 30% sucrose (in PBS) solution for 3 days until brains were fully saturated. Brains were frozen and cryosectioned coronally at 40mm on a sliding microtome. Tissue was stored in a 50% glycerol/PBS solution at −20°C.

### Immunofluorescence

Forty-μm thick free-floating sections were washed in 0.01M tris-buffered solution (TBS; pH 7.4) 3 X for 5 minutes. The tissue then underwent an antigen retrieval process for 30 minutes at 37°C. After antigen retrieval the free-floating sections were washed and blocked in 5% normal serum for 1 to 2 hours. The sections were thereafter incubated in 1% serum in TBS-Triton (TBST) primary antibody solution consisting of one of the following antibodies: anti-Iba1 (1:500, WACO), anti-Olig2 (1:250, clone SP07-02; R&D), anti-GFAP (1:500, clone DIF48; Abcam), anti-CD (1:500, clone 4SM15; eBioscience), anti-Thy1.1 (1:500, OX-7; Invitrogen) anti-CD4 (1:500, clone RM4-5; Thermo Fisher), anti-NK1.1 (1:250, clone EPR22990-12; Abcam). After an overnight incubation, the free-floating sections were washed and put into a 1% serum TBST secondary solution for 2 hours. Sections were mounted onto coated glass slides, and cover slipped using hard set mounting medium (Vector Laboratories). Fluorescence images were collected on a Ti2 Nikon microscope using a Ci2 confocal system.

### DAB labeling and quantification

Free floating striatal sections were quenched in a 3% hydrogen peroxide/50% methanol in 0.01M TBS (pH 7.4) solution at room temp for 5 minutes. After three TBS washes, tissue was incubated in antigen retrieval sodium citrate solution for 30 minutes at 37°C. Background staining was blocked in 5% serum and incubated with either an anti-MHCII (1:500, M5/114.15.2; Thermo Fisher) or a pSer129 (1:5000, clone EPI53644; Abcam) antibody 1% serum overnight at 4oC. The following day, sections were incubated with a biotinylated goat anti-rat IgG secondary antibody (1:1000, Vector Labs) in a 1% serum TBST solution. The R.T.U Vectastain ABC Reagent kit and DAB kits (Vector Labs) were used to develop the stain according to manufacturer’s protocol. Striatal sections were mounted onto plus coated slides and dehydrated with a gradient of ethanol solutions. Lastly slides were cover slipped with Permount (Electron Microscopy Sciences). Slides were imaged at 10X on a Zeiss imager M2 brightfield microscope (MBF Biosciences). Data was analyzed in ImageJ and the fold change of signal between the ipsi- and contra-lateral side were calculated.

### Interferon gamma neutralizing antibody treatment

C57BL/6 mice were pretreated three days before i.p. injection of either a neutralizing IFNγ antibody (clone XMG1.2; 200ng; n=5) or isotype control (IgG1; 200ng; n=5). Three days following the initial i.p. injection, mice received an injection of either Olig001-GFP or Olig001-SYN in the dorsolateral striatum. Immediately after vector injection, mice were given an i.p. dose of their respective treatment. To continue their treatment, mice were injected every three days i.p. with 200ng of either neutralizing IFNγ or the isotype control. After 30 days, mice were anesthetized for their respected endpoints.

### Luxol Fast Blue staining and quantification

Forty-μm thick mounted brain sections were quickly washed with DI water and incubated in a 0.1% Luxol Fast Blue solution at 60°C for 2 hours. Excess dye was removed with running DI water. To differentiate and visualize the myelin from the rest of the tissue, slides were dipped in a 0.05% Lithium Carbonate solution 2X for 1 minute, followed by three 70% ethanol washes. This differentiation step was repeated until the myelin was stained blue and non-lipid parts of the tissue were clear. Slides were quickly dehydrated and mounted with Permount (Electron Microscopy Sciences). Images were taken at 10X on a Zeiss Axio Imager M2 microscope (MFB Biosciences). Images were quantified with ImageJ. Fold change of contra- to ispi-lateral were calculated based on mean grey value.

### Mononuclear cell sorting and flow cytometry

Four weeks post Olig001 delivery, mice were anesthetized and transcardially perfused with 0.01M PBS pH 7.4. Brain tissue was removed and the striata were dissected. Striatal tissues were triturated and digested with 1 mg/mL Collagenase IV (Sigma) and 20 μg/mL DNAse I (Sigma) diluted in RPMI 1640 with 10% heat inactivated fetal bovine serum, 1% glutamine (Sigma), and 1% Penicillin–Streptomycin (Sigma). After enzyme digestion, samples were filtered through a 70 uM filter and mononuclear cells were separated out using a 30/70% percoll gradient (GE).

For all cell labeling, isolated cells were blocked with anti-Fcy receptor (1:100; BD Biosciences). Cell surfaces were labeled with the following fluorescent-conjugated antibodies against CD45 (clone 30-F11; eBioscience), CD11b (clone M1/70; BioLegend), MHCII (clone M5/114.15.2; BioLegend), Ly6C (clone HK1.4; BioLegend), CD4 (clone GK1.5; BioLegend), CD8a (clone 53.6.7; BioLegend), Thy1.1 (clone HIS51; BD Biosciences), or hCD2 (clone RPA2.10; eBioscience). A fixable viability dye was used to distinguish live cells per manufacturer’s instructions (Fixable Near-IR LIVE/DEAD Stain Kit, Invitrogen).

For intracellular transcription factor labeling, the Foxp3/Transcription Factor Staining Kit (eBioscience) was used accordingly with fluorescent-conjugated antibodies against FOXP3 (clone FJK-16S; eBioscience), T-bet (clone 4B10; BioLegend), GATA2 (clone 16E10A23; BioLegend), RORγt (clone Q31-378; BD Biosciences). An Attune Nxt (Thermo Fisher Scientific) or a BD Symphony flow cytometer (BD Sciences) were used to analyze samples and Flow Joe (Tree Star) software were used for analysis. Mean cell count numbers, percentages, and mean fluorescent intensity (MFI) will be measured with FlowJo software to assess for neuroinflammation.

### Statistical Analysis

All graphs and corresponding statistical tests were generated or performed using Prism software (GraphPad). For Flow cytometry data points were compared across time points/antibody treatment using either an independent factorial ANOVA and a Bonferroni’s post hoc test (with 95% confidence and p<0.05) or unpaired students t-test (with 95% confidence and p<0.05).

### Data availability

The authors affirm that the findings of this manuscript are supported by the data therein. Additional information can be requested from the corresponding author.

## Results

### Genetically deleting IFNγ attenuates neuroinflammation in the Olig001-SYN model of MSA

IFNγ induces MHCII expression on the cell surface of APCs^10^. To determine if Olig001-SYN induced neuroinflammation as a result of induction of IFNγ expression, we utilized mice in which the required transcription factor for IFNγ production, Tbet, was deleted (Tbet -/-). Tbet -/- mice and their WT littermate controls, aged 8-12 weeks, received Olig001-SYN (or GFP as control) in the dorsolateral striatum (Fig. 1a). 4 weeks post vector delivery, using immunohistochemistry and mononuclear cell isolation and flow cytometry, we found that Tbet -/- mice showed a reduction in infiltrating Ly6C+ monocytes (CD11b+, CD45hi, Ly6C+) (Fig. 2b and c). Within the monocyte population there was no difference in MHCII expression (Fig. 1c). In response to α-syn expression, Tbet -/- mice displayed no change in the number of microglia within the striatum (Fig. 1c), however, in the absence of Tbet, a significant reduction in MHCII expression was observed on microglia (CD11b+ CD45lo) via flow cytometry (Fig. 1c). Consistent with our flow cytometry results, utilizing immunohistochemistry, we observed a decrease in MHCII expression in the dorsolateral striatum (Fig. 1e and f) indicating that Tbet expression is needed for IFNγ-mediated induction of MHCII expression in the CNS and peripheral monocyte entry.

**Figure 1:**
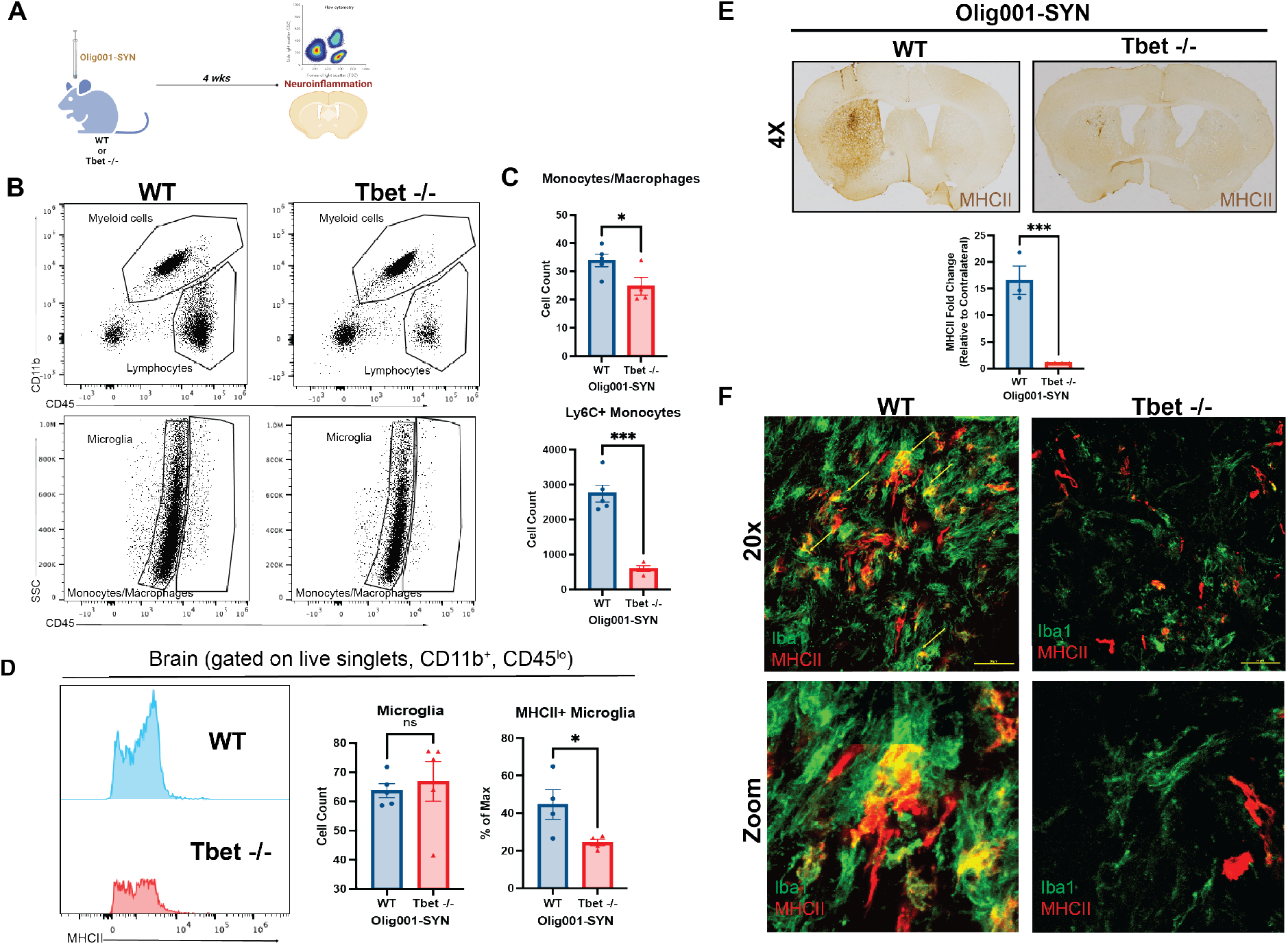
Tbet -/- attenuates myeloid responses in the Olig001-SYN mouse model of MSA. (**A**) Tbet-/- mice or their WT littermates 8-12 weeks of age received bilateral (flow cytometry) or unilateral (immunohistochemistry) stereotaxic injections of Olig001-SYN into the dorsal lateral striatum. 4 weeks post-injection, tissue was collected to assess for neuroinflammation. (**B**) Flow cytometry on isolated striatal tissues, displaying both myeloid (CD45+, CD11b) and lymphocyte populations (CD45+, CD11b-) (top); the resident microglia (CD45lo, CD11b+) and monocytes/macrophages (CD45hi, CD11b+) (bottom). (**C**) Quantification of flow cytometry showing the cell count of MHCII on resident microglia. Mean values are plotted +/- SEM, unpaired t-test, ns = no significance, *p, 0.05. (**D**) The quantification of total monocytes/macrophages (CD45hi, CD11b+) and infiltrating monocytes (CD45hi, CD11b+, Ly6C+) isolated from the striatum with flow cytometry. Mean values are plotted +/- SEM, unpaired t-test, *p < 0.05, ***p < 0005. (**E**) Representative images of 3,3’Diaminobenzidine (DAB) staining and quantification of mean gray value of the MHCII expression in the dorsal lateral striatum. Mean values are plotted +/- SEM, unpaired t-test, ***p < 0005. (**F**) Representative images depicting MHCII expression (red) on activated microglia (Iba1, green) in the dorsal lateral striatum. Scale bars are at 50uM. For immunohistochemistry experiments, n=3 mice per group. For flow cytometry experiments n=3-5 (2 mouse striatum tissues pooled per n) per group.

**Figure 2:**
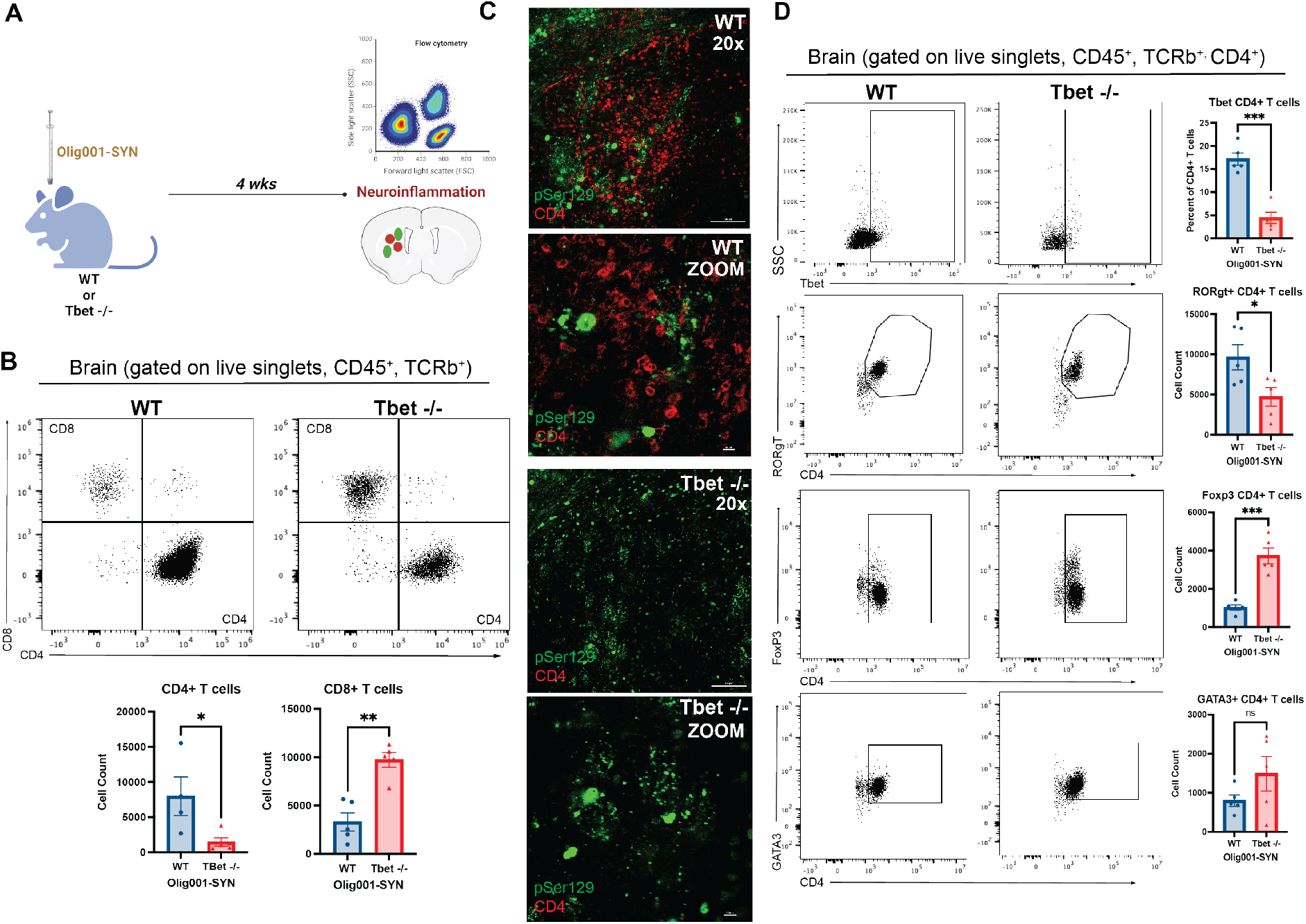
Tbet -/- attenuates CD4+ T cells and alters T cell subsets. (**A**) At 8-12 weeks, Tbet-/- or littermate controls were stereotactically injected with Olig001-SYN into the dorsal lateral striatum. After 4 weeks, tissue was harvested to assess for neuroinflammation. (**B**) Flow cytometry dot plots of CD8+ (CD45+, CD11b-, TCRb+, CD8+) and CD4+ T cells (CD45+, CD11b-, TCRb+, CD4+). Below the dot plots are the cell counts of CD4+ and CD8+ T cells in the striatum in the presence of α-syn pathology within oligodendrocytes. Mean values are plotted +/- SEM, unpaired t-test, *p < 0.05, **p < 001. (**C**) Representative immunohistochemistry images of pSer129 (green) and CD4+ (red) in the dorsal lateral striatum of Tbet -/- mice and their littermate controls. (**D**) Flow cytometry graphs and their corresponding quantification for the following CD4+ T cell subsets: Th1, Th17, T_reg_, Th2. Mean values are plotted +/- SEM, unpaired t-test, ns = no significance, *p< 0.05, ***p<0.005. For immunohistochemistry experiments, n=3 mice per group. For flow cytometry experiments n=5 (2 mouse striatum tissues pooled per n) per group.

To determine whether IFNγ is responsible for T cell infiltration in response to α-syn overexpression, we performed mononuclear cell isolation and flow cytometry 4 weeks post vector delivery in WT and Tbet -/- mice. Tbet -/- mice displayed a decrease in the infiltration of CD4+ T cells (Fig. 2b) and an increase of CD8+ T cells (Fig. 2b) indicating α-syn-mediated neuroinflammation is caused by Th1+ CD4 T cells. Given our previous observations showing that CD4+ T cells are required for MSA pathology, we further investigated the CD4+ T cell response. In the dorsolateral striatum, compared to WT mice, Tbet-/- mice displayed a reduction of CD4+ T cells in close apposition to the pSer129+ GCI pathology (Fig. 2c). Within the CD4+ T cell population, there were significant changes among the T cell subsets Th1, Th17, and T_reg_ including a reduction in the number of Th1 and Th17 cells (Fig. 2d). Conversely, there was a significant increase in the number of T_reg_ within the CD4+ T cell population, suggesting that loss of Tbet mediated IFNγ expression shifts the T cell repertoire from a pro-inflammatory (Th1, Th17) to a restorative (T_reg_) state. As IL-17 produced by T cells can also contribute to proinflammatory or autoimmune responses^19–21^, we performed immunohistochemistry and flow cytometry on WT and RORγt -/- mice treated with both groups were treated with Olig001-SYN. Four weeks post-delivery, RORγt -/- mice overexpressing α-syn in oligodendrocytes showed no attenuation of MHCII expression and enhanced demyelination compared to WT mice (Supplemental Fig 1) indicating that IL-17 producing Th17 cells do not drive Olig001-SYN mediated neuroinflammation and demyelination.

### IFNγ mediates demyelination in the Olig001-SYN mouse model of MSA

Demyelination is a key feature of MSA pathology^1^. To assess how IFNγ contributes to this pathology, Tbet -/- and their WT littermate controls, 8-12 weeks of age, were injected with Olig001-SYN in the dorsal lateral striatum (Fig 3a). At 4 weeks post-transduction myelination was assessed via luxol fast blue staining. Compared to WT mice, Tbet-/- mice displayed a fourfold increase in the degree of myelination in the dorsolateral striatum and corpus callosum (Fig 3c). The preservation of myelin seen in Tbet -/- is comparable to healthy, non-inflamed mouse striatum (Fig 3b) indicating Tbet mediated IFNγ expression is required for demyelination in the Olig001-SYN mouse model.

**Figure 3:**
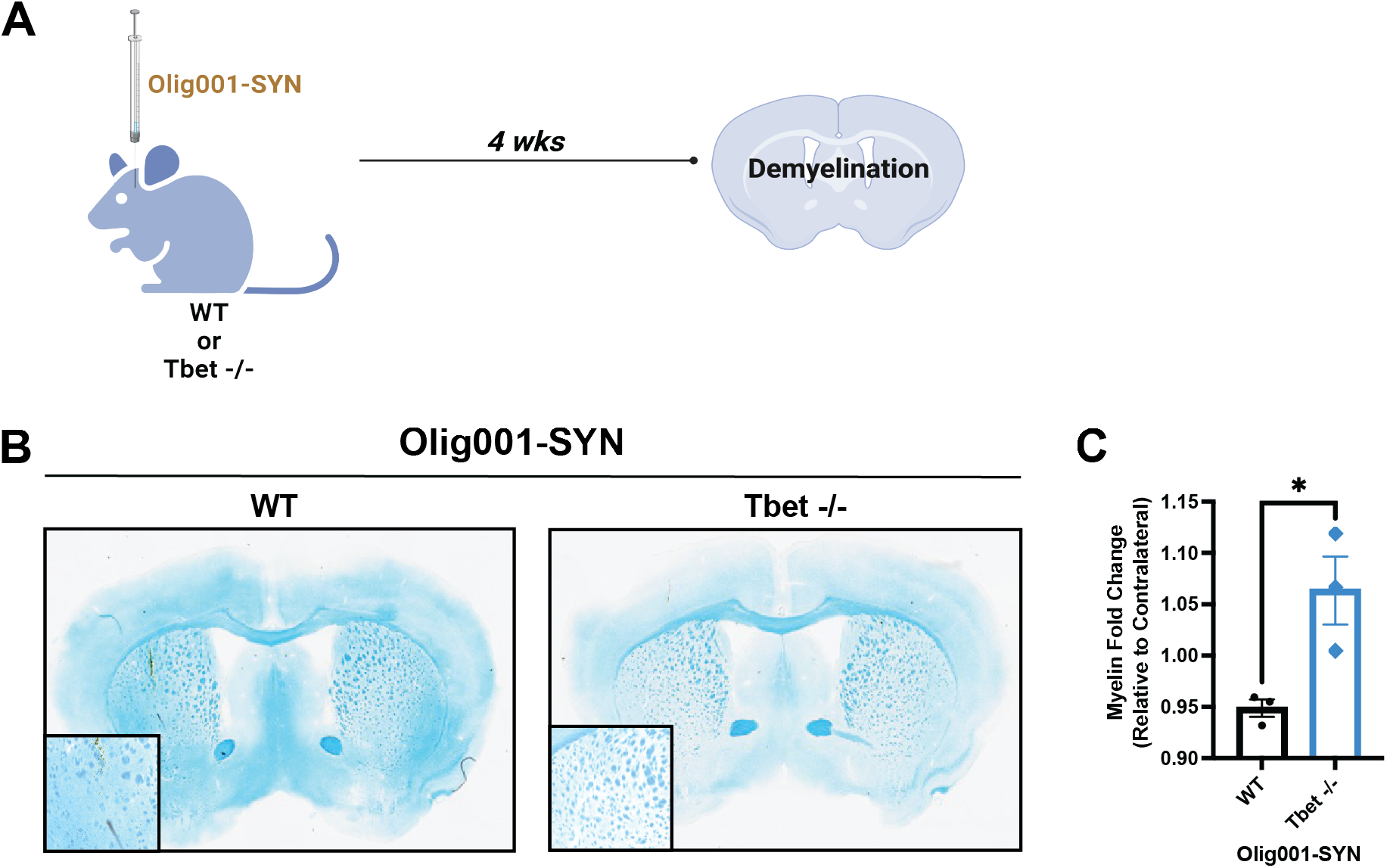
Tbet -/- attenuates demyelination. (**A**) Both male and female Tbet -/- mice and littermate controls were transduced with Olig001-SYN at 8-12 weeks old. 4 weeks post transduction, tissue was collected and stained with Luxol Fast Blue to determine demyelination in the striatum and corpus collosum. (**B**) Representative Luxol Fast Blue images of WT and Tbet -/- where myelinated areas are stained in blue. (**C**) Quantification of the myelin fold change between the ispi- and contralateral sides of the striatum and corpus collosum in WT and Tbet -/- mice. Mean values are plotted +/- SEM, unpaired t-test, *p < 0.05. For immunohistochemistry experiments, n=3 mice per group.

### Pharmacologically targeting IFNγ attenuates Olig001-SYN mediated neuroinflammation

Although the experimentation in the Tbet -/- mouse demonstrated a role of IFNγ in Olig001-SYN-mediated pathology, this holds little translational value. To that end, a key question remains: can targeting IFNγ pharmacologically provide a therapeutic benefit? To determine if IFNγ neutralization attenuates Olig001-SYN mediated neuroinflammation and demyelination, an IFNγ neutralizing antibody was used to globally deplete IFNγ. WT mice (8-12 week old) were pre-treated with either an IFNγ neutralizing antibody (XMG1.2; 200ng) or an isotype control (IgG1; 200ng) intraperitoneally (i.p.). three days prior to Olig001-SYN delivery to the dorsolateral striatum and every three days over the course of 4 weeks. (Fig 4a). Four weeks post-transduction, striatal tissue was harvested to assess for neuroinflammation via immunohistochemistry and flow cytometry.

**Figure 4:**
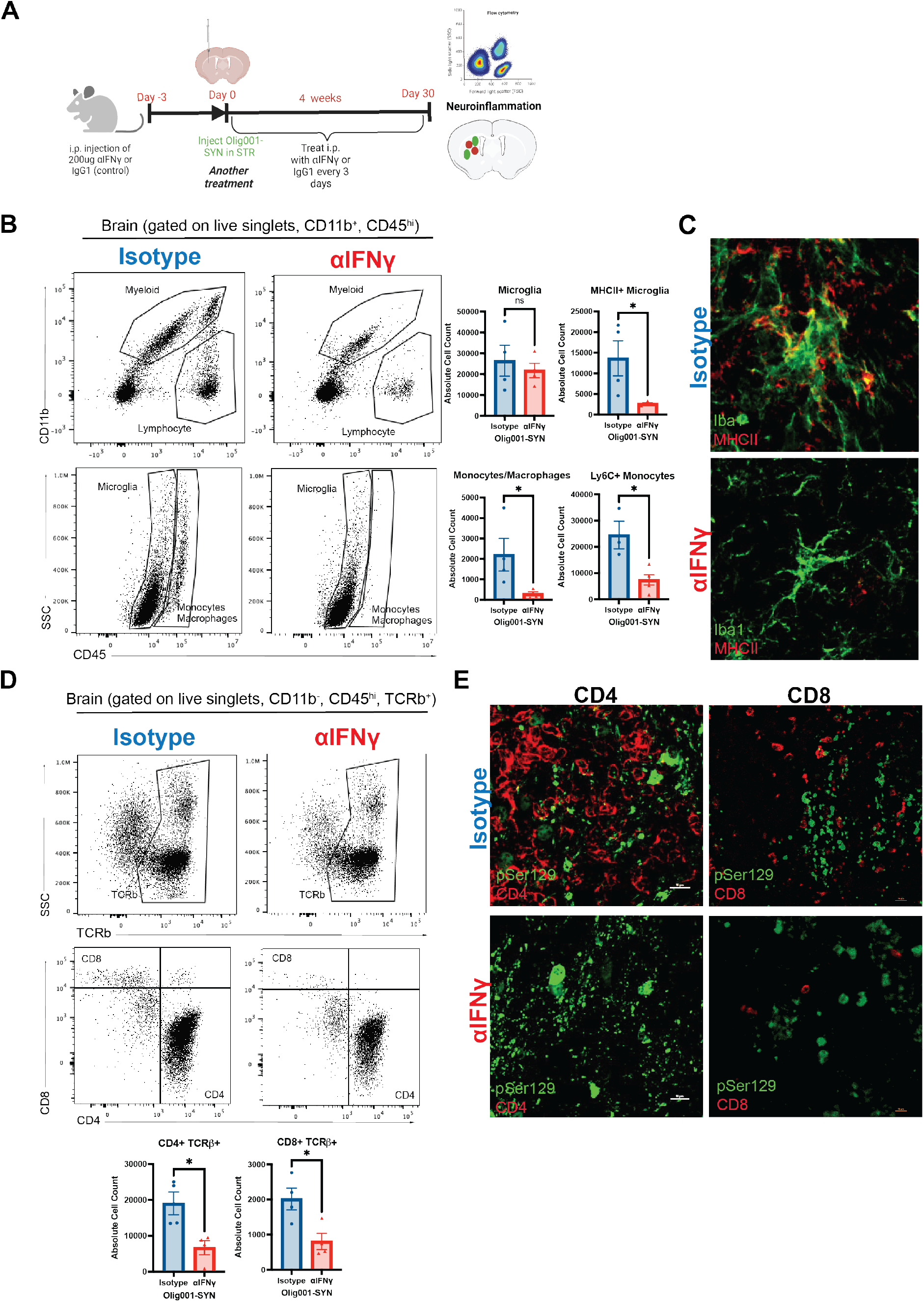
Neutralizing IFNγ attenuates neuroinflammation. (**A**) Three days prior to Olig001-SYN transduction, both WT female and male 8-12 week old mice received an i.p. injection of 200ng of anti-IFNγ (XMG1.2) or its isotype control (IgG1). Three days later, Olig001-SYN was transduced into the striatum and received another treatment. Afterwards mice received follow up treatments every three days for 30 days. After the 30 days, tissue was collected to assess for neuroinflammation. (**B**) Dot plots displaying the overall myeloid populations (CD45+, CD11b+) and lymphocytes (CD45+, CD11b-). Below are dot plots of microglia (CD45^mid^, CD11b+) and monocytes/macrophages (CD45^hi^, CD11b+). Mean values are plotted +/- SEM, unpaired t-test, ns = no significance, *p< 0.05. (**C**) Representative immunohistochemistry of Iba1 (green) and MHCII (red) positive cells within the striatum and corpus collosum. (**D**) flow cytometry of mononuclear TCRb+ (CD45+. CD11b-, TCRb+), CD4+ (CD45+, CD11b-, TCRb+) and CD8+ T cells (CD45+, CD11b-, TCRb+). Mean values are plotted +/- SEM, unpaired t-test, *p< 0.05. (**E**) representative immunohistochemistry images of CD4+ and CD8+ T cells (red) and insoluble alpha-synuclein, pSer129 (green) in the striatum. For immunohistochemistry experiments, n=3 mice per group. For flow cytometry experiments n=4 (2 mouse striatum tissues pooled per n) per group.

Findings show that XMG1.2 treatment significantly attenuated pro-inflammatory monocyte infiltration and MHCII expression on microglia (Fig 4b and c) in the ipsilateral striatum compared to isotype control. Additionally, XMG1.2 treatment significantly decreased the number of CD4+ and CD8+ T cells infiltrating the ipsilateral striatum (Fig 4d). While α-syn expression was unaffected by XMG1.2 treatment (Fig. 4e), peripheral T cell entry was attenuated confirming that IFNγ is mediating α-syn-induced neuroinflammation.

### Pharmacologically targeting IFNγ attenuates Olig001-SYN mediated demyelination

Lastly, to determine if XMG1.2 treatment attenuated Olig001-SYN mediated demyelination, Luxol Fast Blue staining was performed on striatal sections from Olig001-SYN treated mice (Fig. 5a). In this regard, XMG1.2 treatment preserved myelin in the dorsolateral striatum, specifically in the corpus callosum, compared to isotype control (Fig. 5b-c). These results show that neutralizing IFNγ attenuates Olig001-SYN mediated demyelination, again supporting a role of IFNγ expression in Olig001-SYN-mediated demyelination.

**Figure 5:**
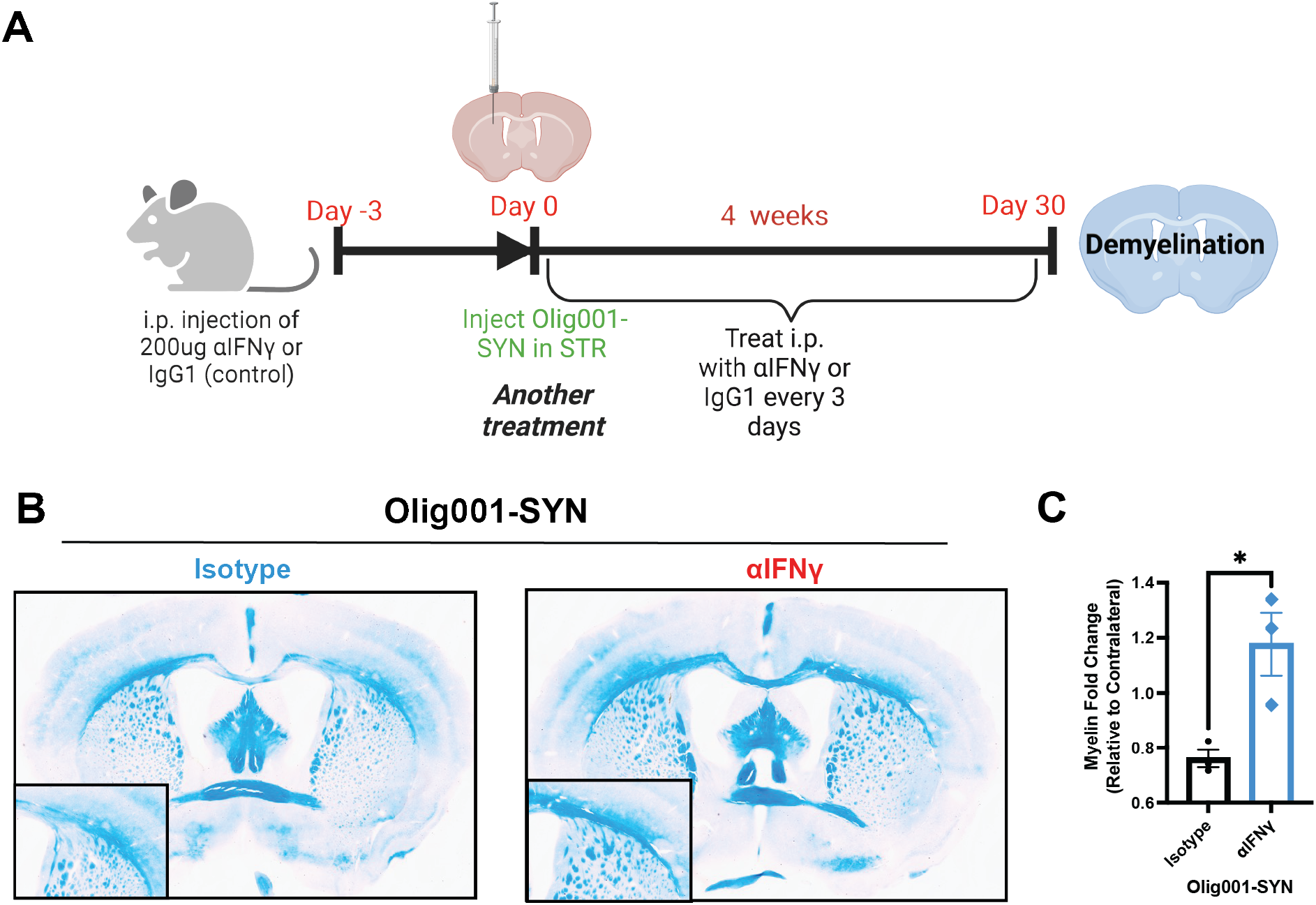
Neutralizing IFNγ attenuates demyelination. (**A**) Same experiment design that was used in assessing neuroinflammation in Figure 4. (**B**) Representative images of Luxol Fast Blue staining displaying myelin within the striatum. Boxes in the bottom left are zooms of areas of demyelination. (C) Myelin fold change between ipsi- and contral-lateral sides of the striatum and corpus collosum. Mean values are plotted +/- SEM, unpaired t-test, *p< 0.05. For immunohistochemistry experiments, n=3 mice per group.

### CD4+ T cells produce IFNγ in response to Olig001-SYN mouse model

In post-mortem MSA tissue there is evidence of infiltrating CD4+ and CD8+ T cells^6^ in the parenchyma and increased IFNγ in the CSF^15^, while studies in preclinical models have shown that CD4+ T cells are required for neuroinflammation and demyelination^6^. Similarly, the results presented here, using genetic and pharmacological approaches, show that IFNγ is a key mediator of the neuroinflammation and demyelination observed in the Olig001-SYN mouse model. While our previous studies in the Olig001-SYN model suggest IFNγ is produced by T cells^6^, it is unclear if other CNS resident and infiltrating immune cells produce IFNγ in response to Olig001-SYN expression in oligodendrocytes. To determine which CNS resident and infiltrating immune cells express IFNγ as a result of α-syn overexpression, we used immunohistochemistry, flow cytometry, and a Thy1.1/IFNγ reporter mouse where Thy1.1 is expressed from the IFNγ promoter^18^. In this mouse, when IFNγ is expressed, Thy1.1 is expressed on the cell surface. Using this reporter model, we isolated mononuclear cells from the dorsolateral striatum of mice treated with Olig001-SYN or Olig001-GFP control (Fig. 6a). Using immunohistochemistry, immune populations known to produce IFNγ (CD4+ T cells, CD8+ T cells, NK cells, astrocytes, and microglia) were investigated for Thy1.1 expression (Fig 6b). These lymphocytes (CD45+) were analyzed by flow cytometry and our results show that CD4+ T cells expressed the overwhelming majority of Thy1.1 on the cell surface in response to α-syn overexpression in oligodendrocytes (Fig. 6c) which was significantly elevated as compared to the GFP control. Upon further investigation, the majority of Thy1.1 expressed was found on CD45+ TCRb+ CD4+ T cells (Fig 6d-f), matching results seen in the immunohistochemistry (Fig 6b). Given the CD4+ T cells are producing the pathogenic IFNγ (Fig 6f) when α-syn is expressed in oligodendrocytes, these data suggest that the CD4+ T cell subtype, Th1 cells are facilitating the disease process via production of IFNγ. In summary, our results show other immune cell types like CD8+ T cells, B cells, and NK cells do not contribute to significant expression of IFNγ, but CD4+ T cells drive MSA pathology via IFNγ expression.

**Figure 6:**
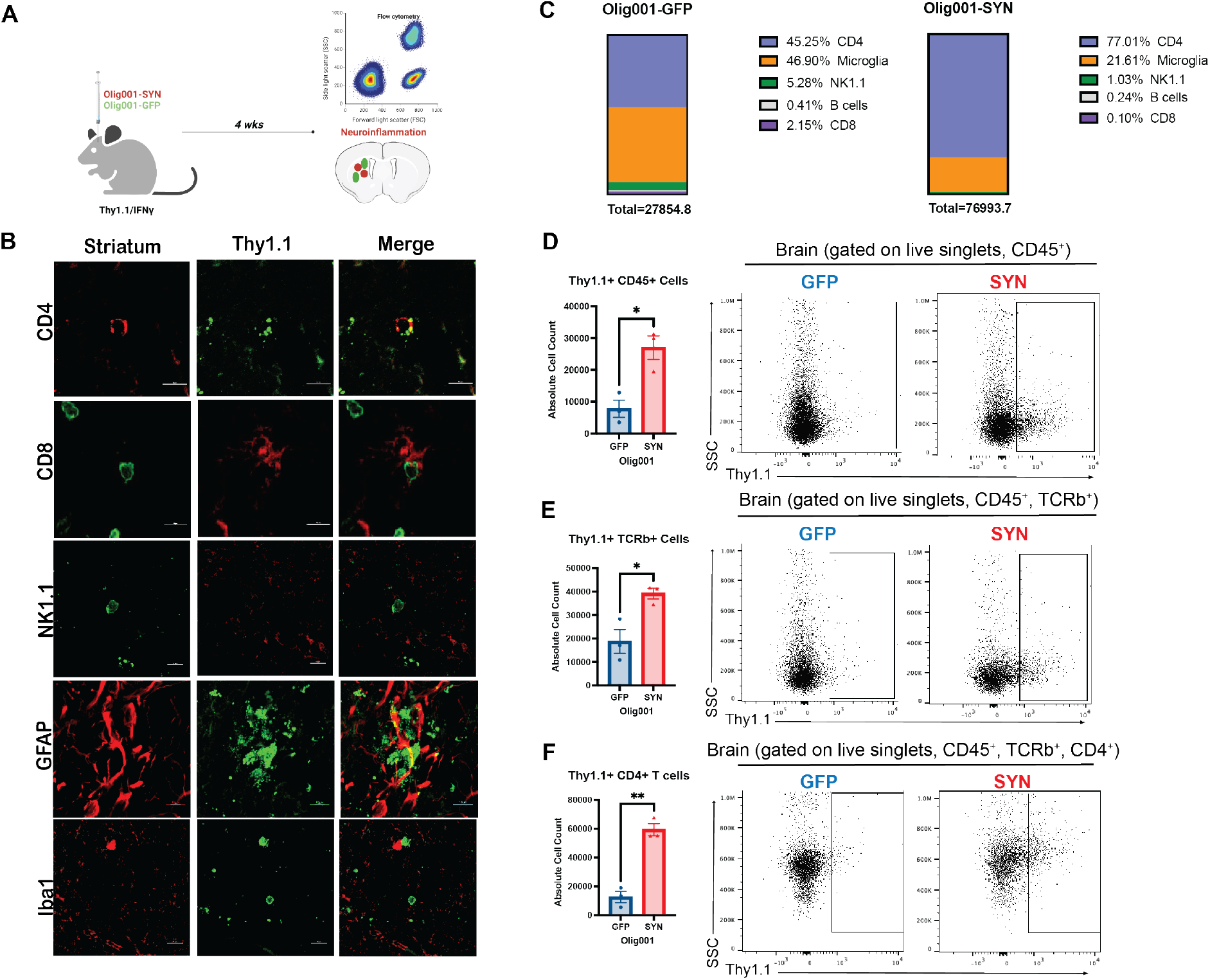
Proinflammatory IFNγ is produced by CD4+ T cells. (**A**) Male and female 8–12- week-old Thy1.1/IFNγ reporter mice were injected with either Olig001-GFP or Olig001-SYN in the dorsal lateral striatum. 4 weeks post-transduction, tissue was harvested for neuroinflammation. (**B**) A panel of representative immunohistochemistry images of CD4+ T cells (CD4), CD8+ T cells (CD8), NK cells (NK1.1), astrocytes (GFAP), and microglia (Iba1) with the IFNγ reporter Thy1.1. (**C**) Percentages of CD45+ cells generated from flow cytometry data looking at Thy1.1+ (IFNγ producing) cells. (**D**) Dot plots between Olig001-GFP and Olig001-SYN injected Thy1.1/IFNγ reporter mice showing Thy1.1+ cells in lymphocytes (CD45+, CD11b-). Mean values are plotted +/- SEM, unpaired t-test, *p< 0.05. (**E**) Flow cytometry dot plots of Thy1.1+ TCRb+ T cells (CD45+, CD11b-, TCRb+, Thy1.1+). Mean values are plotted +/- SEM, unpaired t-test, *p< 0.05. (**F**) Flow cytometry showing Thy1.1+ CD4+ T cells (CD45+, CD11b-, TCRb+, CD4+, Thy1.1+). Mean values are plotted +/- SEM, unpaired t-test, *p< 0.05. For immunohistochemistry experiments, n=3 mice per group. For flow cytometry experiments n=3 (2 mouse striatum tissues pooled per n) per group.

## Discussion

In this study we show that IFNγ, primarily produced by CD4+ T cells, induces neuroinflammation and demyelination in the Olig001-SYN mouse model of MSA. Using a genetic approach in which IFNγ was globally knocked out, neuroinflammation and demyelination were attenuated four weeks post vector delivery. Additionally, when IFNγ was pharmacology depleted with the IFNγ neutralizing antibody (XMG 1.2), there was a significant decrease in activated CNS myeloid populations, MHCII expression, and infiltrating CD4+ and CD8+ T cells. Lastly, using a novel IFNγ reporter mouse, we found that the Olig001-SYN mediated increase in IFNγ expression originates from CD4+ T cells, suggesting that Th1, not Th17 T cells, are key in facilitating neuroinflammation and demyelination. Overall, our results indicate that IFNγ mediates the neuroinflammation and demyelination in this mouse model of MSA, and thus targeting IFNγ producing CD4+ T cells may be a future disease modifying therapeutic.

Although there has been extensive research into other demyelinating disease like MS, the role alpha-synuclein in oligodendrocytes play is still unclear^22^. Our results show that blocking IFNγ prevents demyelination (Fig 1-5). Under healthy conditions, cytokines and chemokines can affect oligodendroglial differentiation^23–25^. For example, IFNγ stimulates oligodendroglia to present MHCI and MHCII mediated antigens in the context of MS^23,26^. Oligodendrocytes are responsive to IFNγ in MSA, still, the mechanism of how oligodendrocytes contribute to disease pathogenesis is unknown. In MSA, α-syn aggregation contributes to demyelination and inflammation, however, it is unclear if demyelination occurs in a cell autonomous or non-cell autonomous manner^27^. It has been suggested that disease-associated oligodendrocytes participate in crosstalk with astrocytes and microglia to help mediate inflammatory responses^25^. While there is growing evidence for disease associated oligodendrocytes in MS and Alzheimer’s disease (AD)^25^, it is currently unknown if there are disease-associated oligodendrocytes and if they communicate with other glia and other immune populations in synucleinopathies such as MSA. Future studies are needed to understand the mechanisms behind α-syn mediated demyelination, and if oligodendrocytes mediate neuroinflammatory responses in MSA.

In this study, Tbet deficiency resulted in significant attenuation of CD4+ T cell infiltration, CNS myeloid activation, and infiltrating monocytes, indicating that genetically deleting IFNγ attenuates Olig001-SYN mediated neuroinflammation (Fig 1-2). While genetic deletion of Tbet attenuated CD4 T cell infiltration and shifted the T cell repertoire to disease resolving, interestingly, there was a significant increase in CD8+ T cells. Although unexpected, this observation highlights either a compensatory effect of Tbet deficiency or a possible role of CD8+ T_regs_ in repressing an inflammatory response. Growing evidence in human and mouse MS studies highlight the existence of CD8+ T_regs_, and their role in suppressing proinflammatory cells like Th1 and Th17^19,20,28^. In human post-mortem MSA tissue, our previous study showed a significant presence of CD8+ T cells^6^. However, studies in blood have shown a decrease in CD8+ T cells and a significant shift in the CD4/CD8 ratio favoring CD4+ T cells^29^. While our previous studies have shown CD4+ T cells are important in disease progression, there is little evidence in the literature investigating the role of CD8+ T cells in MSA. Further studies are needed to understand the role of CD8+ T cells in MSA and whether they mediate neuroinflammatory and neurodegenerative responses via their cytotoxic effector functions.

Using a novel Thy1.1/IFNγ reporter mouse model, we were able to show that the majority of Olig001-SYN mediated IFNγ production originates from infiltrating CD4+ T cells. This, combined with the observation that Tbet -/- mice and CD4 -/- mice show attenuated neuroinflammation and demyelination (Fig 1-3), strongly suggests that Th1 cells are pushing disease progression in MSA. Although our data suggests that Th1 cells are the main driver of MSA pathology, they are not the only CD4+ T cell subtype known to be associated with inflammatory or autoimmune disorders^19–21,30^. Th17 cells have been shown to induce demyelination and inflammation in MS^19,20,30^ via production of the proinflammatory cytokine IL-17a^21^. While increases in Th17 and IL-17a were observed in the Olig001-SYN model in previous studies, using a RORγt -/- mouse, we were able to show that this pathogenic subtype does not cause the neuroinflammation and demyelination in this model, again supporting the notion that Th17 cells are not key to disease progression (Supplemental Fig 1). Future studies in human post-mortem brain and blood are warranted to determine the role of Th1 and Th17 cells in MSA pathogenesis.

In addition to genetic knockout studies, we also showed that neutralizing IFNγ via XMG1.2 treatment was effective in attenuating the neuroinflammation and demyelination in the Olig001-SYN model (Figs 3-5). This observation not only highlighted the importance of IFNγ in facilitating MSA pathology, but it also identifies a potential disease-modifying therapeutic modality in MSA. Currently, there are no disease modifying treatments for MSA^1,22^, only limited symptomatic treatments. The symptomatic treatments do not extend the lifespan of MSA patients, as they do not halt or slow disease progression. Current clinical trials are designed to target the accumulation of α-syn pathology, or active/passive immunization^22,31^. Although early results from preclinical models were promising, most trials were terminated in early phases due to failure to meet primary endpoints or the clinical trial sizes were extremely small. While results in animal models suggest targeting neuroinflammation would be promising^5,32^, new clinical trials have targeted the robust neuroinflammatory response observed in MSA. Approaches to suppress activated microglia or astrogliosis failed in phase II due to patients still presenting with neuroinflammation and rapid clinical decline indicating that a peripheral cell type may be driving disease pathogenesis^22,31^. Intravenous immunoglobin (IVIG) therapy targeting reactive T cells has shown promising phase II results, however, due to the small trial size, this intervention needs further study and development. The current study identifies the IFNγ pathway as a potential disease modifying therapeutic target in a novel mouse model of MSA. Future studies are needed to determine the timing for optimal therapeutic benefit and to determine whether targeting IFNγ attenuates neurodegeneration in the human disease.

In conclusion, the results from these studies show that IFNγ is a key facilitator of neuroinflammation and demyelination as a result of α-syn overexpression in the Olig001-SYN mouse model of MSA. Specifically, genetic knockout or pharmacological approaches targeting IFNγ expression or signaling attenuated CNS microglial activation and infiltration of pro-inflammatory monocytes and CD4+ T cells. Additionally, targeting IFNγ expression or signaling attenuated Olig001-SYN mediated loss of oligodendrocytes and demyelination. Using a novel Thy1.1/IFNγ reporter mouse, we determined that IFNγ was expressed primarily by infiltrating CD4+ T cells in response to α-syn overexpression, suggesting Th1 CD4 T cells are key in facilitating inflammation and disease progression. These findings indicate that IFNγ represents a potential future disease-modifying therapeutic target in MSA.

## Abbreviations

α-syn: alpha-synuclein
APC: antigen presenting cells
GCI: glial cytoplasmic inclusions
IFNγ: Interferon gamma
IFNγR1: interferon gamma receptor 1
MHCII: major histocompatibility complex II
MSA: multiple system atrophy
PD: Parkinson’s disease

## Acknowledgements

The work in these studies was generously supported by NIH/NINDS R01NS107316 (Harms, PI). The authors are grateful for the donation of the Thy1.1/IFNg and hcd2/IL-17a reporter provided by Casey Weaver. We are thankful for all the undergraduates, especially Cameron Gallups, who assisted in various experiments. We are also grateful for the experimental input, discussion, and key reagents provided by Chander Raman. We are also thankful for Vidya Sagar Hanumanthu and the UAB Comprehensive Flow Cytometry Core for their assistance in flow cytometry experiments. Experimental design figures were generated with BioRender.

