## Supplemental Figure 1 for "IFNγ drives neuroinflammation and demyelination in a mouse model of multiple system atrophy"

### Supplementary material

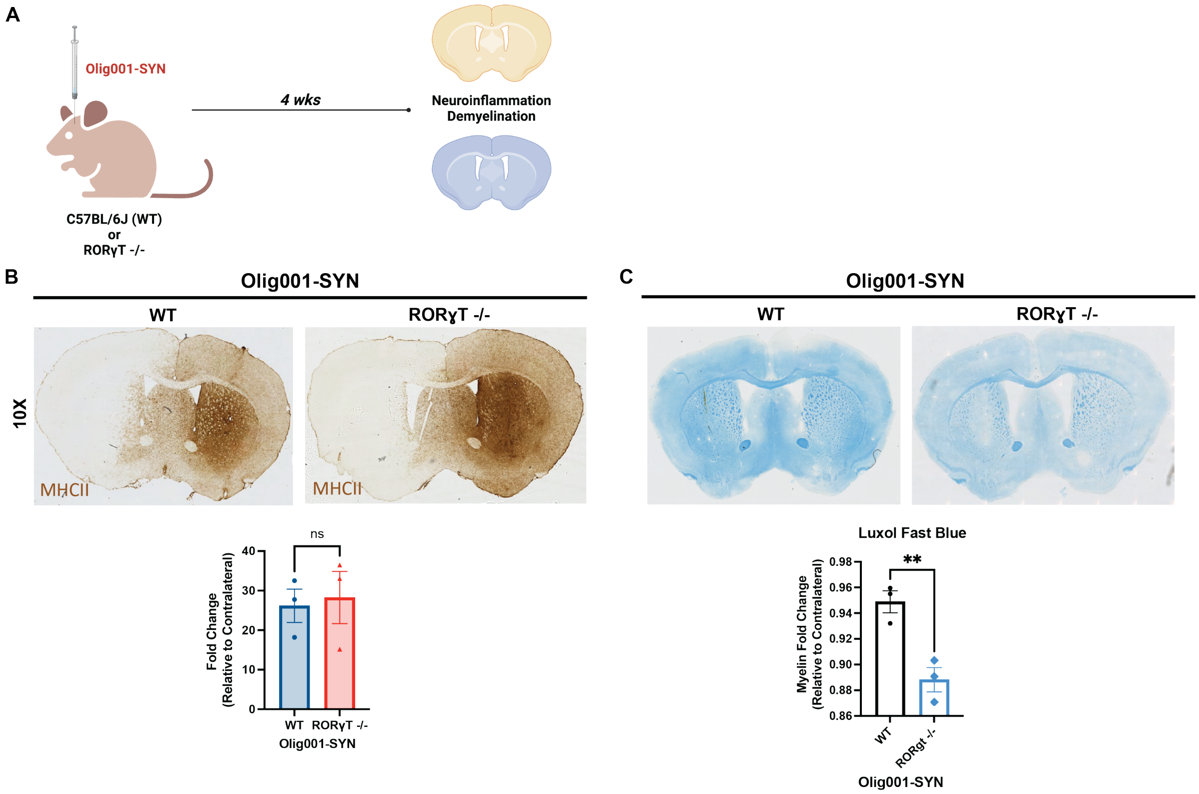


**Supplementary Figure 1: RORγT -/- mice show no attenuation of MHCII expression and enhanced demyelination.** (**A**) Both male and female RORγT -/- mice and littermate controls were transduced with Olig001-SYN at 8-12 weeks old. 4 weeks post transduction, tissue was collected and stained with DAB and Luxol Fast Blue to determine demyelination in the striatum and corpus collosum. (**B**) Representative DAB images of WT and RORγT -/- mice where MHCII staining is highlighted by staining in the striatum. Quantification between the ispi- and contralateral sides of the striatum in WT and RORγT -/- mice were calculated using fold change of staining intensity relative to the contralateral side of the brain. (C) Representative images of Luxol Fast Blue staining of WT and RORγT -/- mice. Below is the quantification of the fold change between the ispi- and contralateral sides. Mean values are plotted +/- SEM, unpaired t-test, **$p < 0.001$. For immunohistochemistry experiments, n=3 mice per group.
